# Bettering of Learning Activity through Elevated P-Cresol Levels in the Brain: Insights from Active Avoidance Testing in Wistar Rats

**DOI:** 10.1101/2024.03.25.586517

**Authors:** Gigi Tevzadze, Zaal Kikvidze

## Abstract

It is established that p-cresol, a compound produced by bacterial colonies within the gastrointestinal tract of mammals, plays a contributory role in the manifestation of various mental disorders. Recently, our research demonstrated that diminishing p-cresol levels in the brain adversely impact the behavioral manifestation of cognitive abilities in rats. In this study, we aimed to investigate the impact of augmenting p-cresol levels in the brain on learning. The Active Avoidance Test was employed to assess learning capabilities. The results, with a high level of confidence, indicated that rats with an increased concentration of p-cresol in the brain exhibited superior task performance and accelerated learning compared to the control group.

## Introduction

P-cresol, a byproduct of bacterial fermentation in mammalian intestines, has been implicated in various mental disorders. Several studies on children’s groups in different countries show a positive, high-confidence correlation between mental disorders, such as autism and epilepsy, with elevated amounts of p-cresol in the urine (Altieri, 2011; Gabriele, 2014; Tevzadze, 2017); Moreover, several animal experiments show, that elevated amount of p-cresol can induce depression-like disorders, and heightened epilepsy reactions (Tevzadze, 2018; Tevzadze, 2018^1^). In recent years, more and more studies have linked gut microbiota with different mental incapabilities, such as Alzheimer’s and Parkinson’s diseases (Bostanciklioglu, 2018; Zhu, 2022). However, the role of p-cresol and gut microbiota should not be reduced only to altering mental health: Recent evidence also suggests a potential link between gut microbiota, specifically *Clostridium difficile* levels, and cognitive activity (Tevzadze, 2023). Building upon these findings, this study explores the impact of gut bacteria-produced compounds, specifically p-cresol, on cognitive activity, particularly in the context of the learning process.

## Material and Methods

### Animals

Animal care during the experimental procedures was carried out by the recommendation of the Ilia State University Research Projects Ethics Commission (Decision no. R/429-23) and by the Council of Europe Directive 2010/63/EU for animal experiments. A total of 14 male young (10 weeks old) rats (weight, 150-180 g) male Wistar rats were obtained from the breeding colony of the vivarium of I. Beritashvili Center of Experimental Biomedicine (Tbilisi, Georgia), were housed in cages (7 rats per cage) and provided with food and water available *ad libitum* and maintained under conditions at a temperature of 20-22 [C and 40-55% humidity on a 12-light/dark cycle (lights on at 7:00 a.m.). The rats were subjected to an enforced swimming test in order to exclude endogenic depression.

Prior to commencing treatment, all rats were marked and separated for 24 h.

### Treatment with p-cresol

After testing rats were randomly assigned into the following subgroups: I) The control group (n=7); and II) The p-cresol-treated group (n=7).

P-cresol treatment initiated after a 3-day acclimatization period. The amount of p-cresol and timing is defined according to previous experiments (Tevzadze, 2018^1^) to avoid depression-like behavior induced through the administration of high doses of p-cresol.

P-cresol (20 mg/kg/day) in 1% ethanol saline solution was administered intraperitoneally during 10 days, whereas control group were subjected for the 1% ethanol saline injection (placebo).

### Active avoidance test

Active avoidance is a classical conditioning test in which rodents must pair the presence of a conditioned stimulus with moving between two chambers to avoid an electric shock (unconditional stimulus). In the active avoidance test, the test animal learns to avoid an aversive stimulus by changing locations. At the start of the test, a rat is placed in one of two compartments. After habituation, a stimulus (light) is presented for a fixed period of time and followed by electrical stimulation of the paws. The rat learns to avoid the shock by moving into the adjacent compartment upon the appearance of the conditioned stimulus (Jänicke, Coper, 1996; Choi, 2010). the faster the rat learns the avoidance, the better its learning ability and the number of avoidance while pairing. In total, each rat in both groups received 25 pairings.

On the second day after the last treatment, the active avoidance test (AAT) was conducted in operant chambers (12.0’’ L x 9.5’’ W x 8.25’’) inside sound-attenuating cabinets. Chambers were fitted with footshock grids, retractable levers, cue lights above the levers, a house light, and speakers for cue tone and white noise.

‘‘Escape’’ and ‘‘avoidance’’ responses were tallied and data are presented as the percentage of trials on which rats emitted an avoidance response [(# of avoidance responses per session/ total number of responses in session) *100].

### Statistical Analysis

We used the paired samples t-test to compare the means of two samples (control versus p-cresol-administered) since each observation in the control sample could be paired with an observation in the experimental sample. As a complementary analytical tool, we used the logistic curve model — a popular approach when analyzing learning curves in animal groups (Gallistel et al. 2004). For our case, in which the learning curve shows the change in the number of avoidance reactions as dependent on the number of trials, the model can be presented as follows:

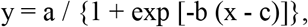

where y is the learned avoidance rate and x is the number of trials, a is the maximum rate of the learned avoidance reached at the sufficiently large number of trials, b is the steepness of the curve and c represents the inflection point, the value of x at which the curve growth turn accelerated to decelerated. The model fitting was assessed by the Akaike Information Criterion (AIC) and the determination coefficient R^2^. Both characteristics showed an acceptable to good accuracy of model fitting: AIC < 7.56, R^2^> 0.99. The paired test was performed, the logistic models have been constructed and their parameters calculated using the software PAST 4 (Hammer and Harper, 2001).

### Results

The p-cresol-administered rats consistently reached higher rates of avoidance as compared to the control group (t = -6.446, p < 0.0001 by the paired t-test, Figure 1). Generally, the learning curve in the p-cresol-treated rats was steeper and would reach a higher maximum rate of avoidance with increasing numbers of trials (Table 1, Figure 1). In particular, the model fitting was accurate both for the control and experimental curves (AIC = 7.46, R^2^= 0.997 and AIC = 7.57, R^2^= 0.998, respectively), and showed that the inflexion point (parameter c) was the same for both curves, meaning that the rats of both groups could reach their maximum rates of avoidance simultaneously; however, because p-cresol-administered rats reached higher maximum rate of avoidance (28% increase in parameter a), their learning curve was steeper (56% decrease in parameter b).

**Figure 1:**
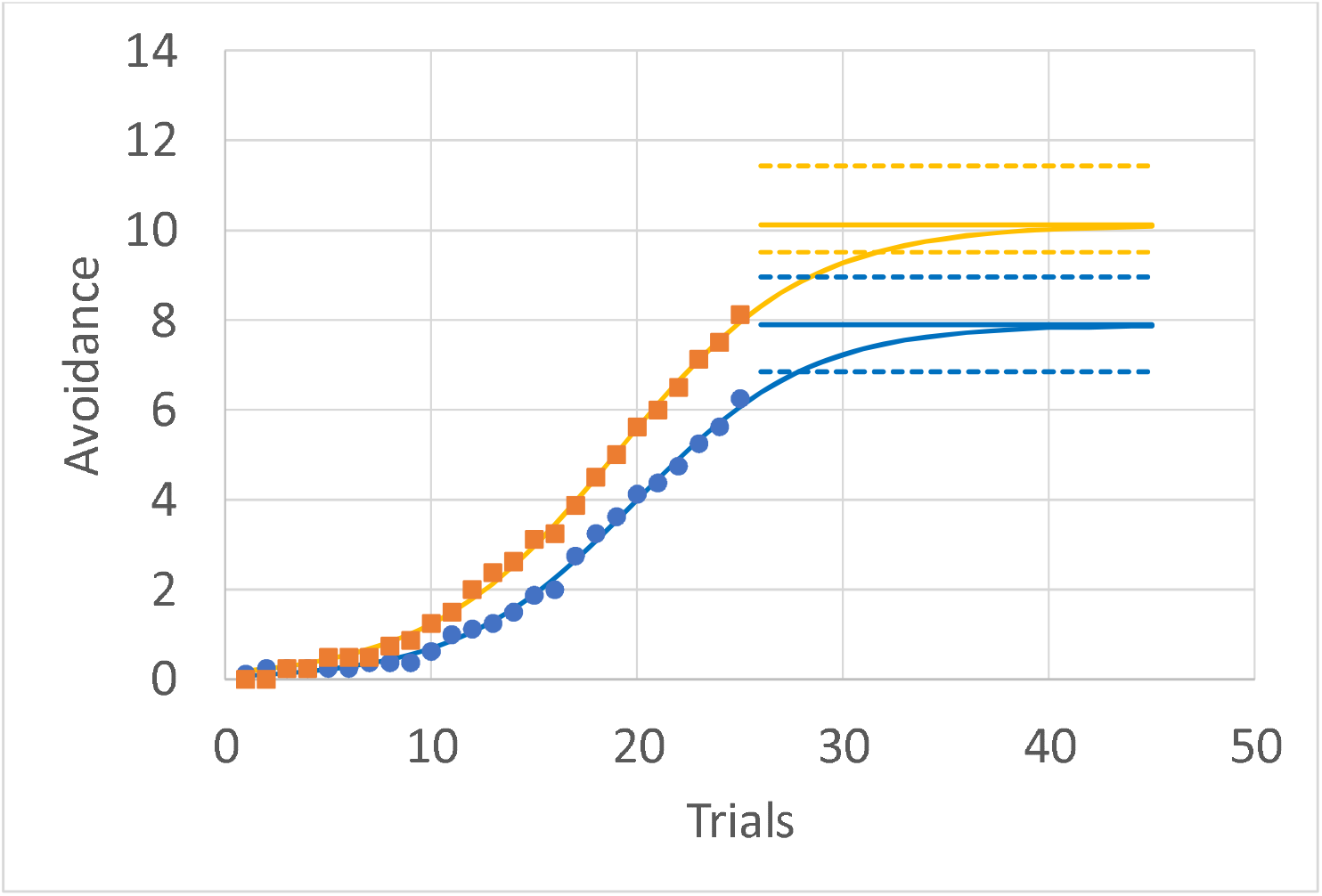
The avoidance learning curve in the control (blue lines and circles) and p-cresol administered (orange lines and squares) young Wistar rats The avoidance learning curve in the control (blue lines and circles) and p-cresol administered (orange lines and squares) young Wistar rats. Circles and squares are the observed data, the curves of the same color show corresponding calculated values, solid horizontal lines are asymptotes. The dashed lines above and below the asymptotes show high and low 95% confidence intervals for each asymptote. These intervals do not overlap to show that the difference in the asymptotes between the control and experimental groups is statistically significant (see also Table 1).

**Table 1.**
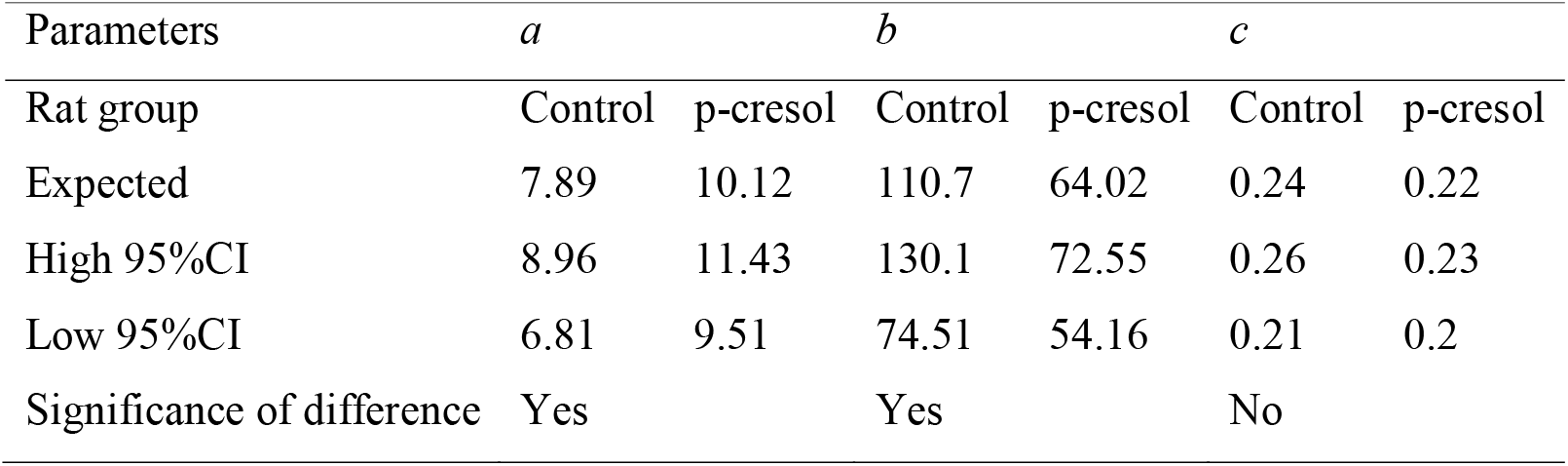
The parameters of the learning curve model (logistic curve *y* = *a* / {1 + exp[*b*(*x -c* )]}. The expected values and their 95% confidence intervals (95%CI) are shown.

| Parameters | *a* |  | *b* |  | *c* |  |
| --- | --- | --- | --- | --- | --- | --- |
| Rat group | Control | p-cresol | Control | p-cresol | Control | p-cresol |
| Expected | 7.89 | 10.12 | 110.7 | 64.02 | 0.24 | 0.22 |
| High 95%CI | 8.96 | 11.43 | 130.1 | 72.55 | 0.26 | 0.23 |
| Low 95%CI | 6.81 | 9.51 | 74.51 | 54.16 | 0.21 | 0.2 |
| Significance of difference | Yes |  | Yes |  | No |  |

## Discussion

Our results clearly show that rats in the p-cresol-treated group exhibited enhanced learning ability compared to the control group. Statistically significant increase was observed with all p-cresol-treated rats achieving higher rate of learned avoidance (expressed in the increasing number of avoidances during pairing sessions) as compared to their control counterparts after the similar number of trials.

Our findings are consistent with our prior investigations (Tevzadze, 2021), which revealed that p-cresol modulates dopaminergic systems and triggers the release of brain-derived neurotrophic factor (BDNF) in nerve cells. Through study, the impact of p-cresol on BDNF secretion and neurofilament subunit expression was explored using rat pheochromocytoma cells (PC-12 cells). The research observed that specific p-cresol doses potentiated nerve growth factor-induced differentiation through the secretion of BDNF in cultured PC-12 cells.

Furthermore, our observations align with our earlier work (Tevzadze, 2020; Tevzadze 2019), indicating the influence of p-cresol on N-methyl-D-aspartate (NMDA) receptors. Specifically, it was noted that the content of the NR2B subunit of NMDA glutamate receptors increased in rats treated with p-cresol.

In our recent experiment (Tevzadze, 2023), a reduction in p-cresol levels, achieved by decreasing the population of *Clostridium difficile* (the primary producer of p-cresol in gut microbiota) resulted in a negative impact on cognitive activity in rats. This interaction between gut microbiota and the brain was proposed to be mediated through the dopaminergic activity of the latter.

The outcomes of the Active Avoidance Test (AAT) suggest that the activity of p-cresol in the prefrontal lobe, responsible for attention, mirrors its effects observed in PC-12 cells and on NMDA receptors. Thus, by activating BDNF and inducing oxidative stress, p-cresol appears to enhance the attention process, offering a potential explanation for the superior learning performance observed in rats treated with p-cresol in our conducted experiment.

## Acknowledgements

The authors express their gratitude to Ms. Tamar Shetekauri and Mr. Nino Tkemaladze for their valuable assistance in conducting AAT in the laboratory.

## Availability of data and materials

The datasets used and/or analyzed during the current study are available from the corresponding author upon reasonable request.

## Authors’ contributions

GT proposed the main hypothesis for the study, designed and supervising the main experiments, and wrote of the manuscript; ZK performed the statistical analysis and prepared figure and table.

## Ethics approval and consent to participate

Animal care during experimental procedures was carried out by the recommendation of the Ilia State University Research Projects Ethics Commission (Decision no. R/266-23) and by Council of Europe Directive 2010/63 / EU for animal experiments.

## Patient consent for publication

Not applicable.

